# A curated dataset of modern and ancient high-coverage shotgun human genomes

**DOI:** 10.1101/2020.10.27.351692

**Authors:** Pierpaolo Maisano Delser, Eppie R. Jones, Anahit Hovhannisyan, Lara Cassidy, Ron Pinhasi, Andrea Manica

## Abstract

Over the last few years, genome-wide data for a large number of ancient human samples have been collected. Whilst datasets of capture SNPs have been collated, high coverage shotgun genomes (which are relatively few but allow certain type of analyses not possible with ascertained captured SNPs) have to be reprocessed by individual groups from raw reads. This task is computationally intensive. Here, we release a dataset including 34 whole-genome sequenced samples, previously published and distributed worldwide, together with the genetic pipeline used to process them. The dataset contains 73,435,604 sites called across 18 ancient and 16 modern individuals and includes sequence data from four previously published ancient samples which we sequenced to higher coverage (10-18x). Such a resource will allow researchers to analyse their new samples with the same genetic pipeline and directly compare them to the reference dataset without re-processing published samples. Moreover, this dataset can be easily expanded to increase the sample distribution both across time and space.

## Background & Summary

The number of ancient humans with genome-wide data available has increased from less than five a decade ago to more than 3,000 thanks to advancements in extraction and sequencing methods for ancient DNA (aDNA)^1^. However, there are just a few high-quality (coverage > 10x) shotgun whole-genome sequenced ancient samples^2^. Moreover, the genetic pipelines used to process shotgun aDNA data are very diverse, making it hard to combine published samples from different studies and research groups. Therefore, researchers have to download raw reads of published samples and reprocess them to create a dataset to compare their new samples against to without pipeline-associated biases. This problem is less pronounced for modern DNA samples as the higher quality of DNA and sequencing coverage partially reduce the biases introduced by the usage of different bioinformatic tools.

Panels including shotgun data for modern samples distributed worldwide have been previously published, such as the Simon Genome Diversity Program^3^, 1000 Genome Project^4^ and Human Genome Diversity Project (HGDP-CEPH panel)^5^.

However, the same concept has not yet been applied to ancient samples or a mix of modern and ancient samples. This study aims to start filling this gap by creating a dataset including both modern and ancient samples distributed across all continents. Therefore, we fully reprocessed 14 high-quality shotgun sequenced ancient samples downloaded from the literature, generated additional new data for previously published 4 ancient samples and merged them with 16 modern samples. The final dataset includes 34 individuals and researchers can use it to quickly compare their new samples against a set of individuals distributed across time and space (Figure 1). Moreover, we hope that researchers will add additional data processed with the pipeline that we released to increase the sample resolution both in time and space.

**Figure 1:**
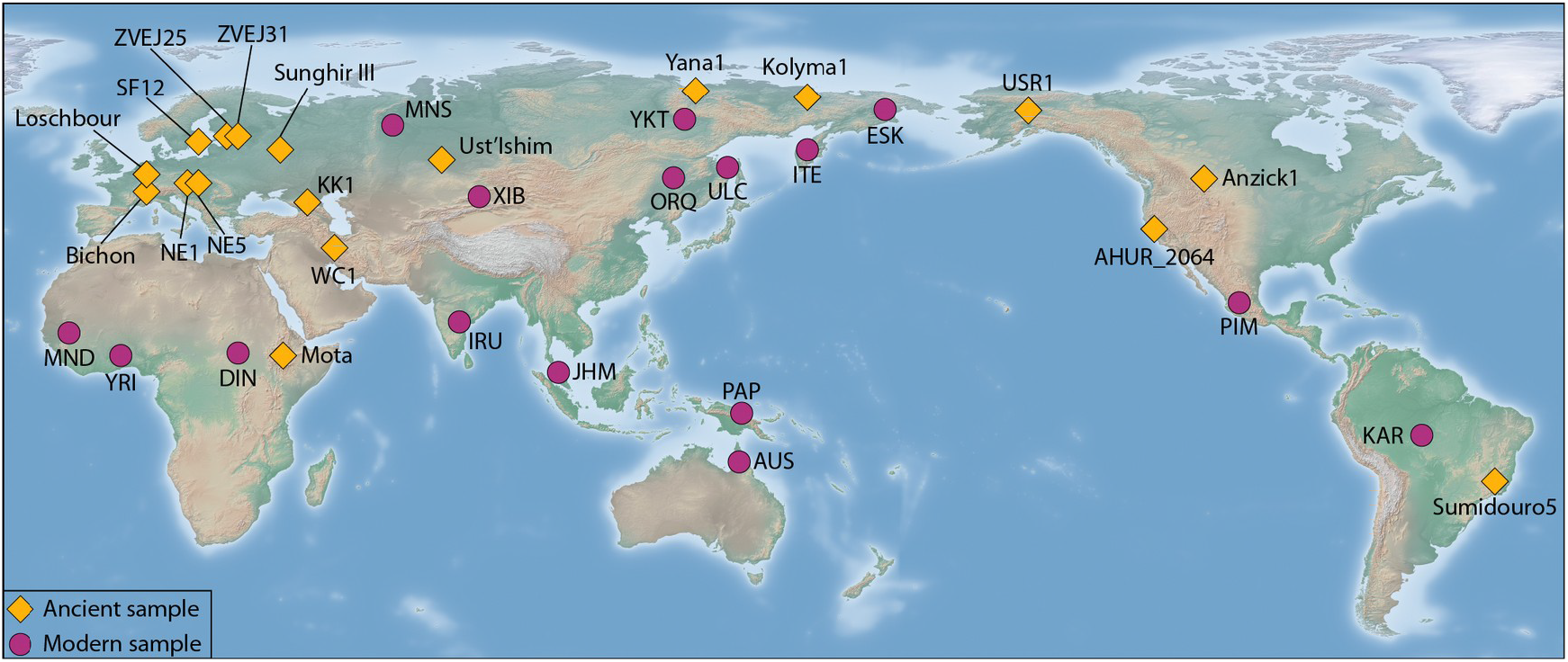
Geographic distribution of samples included in the dataset. Population acronyms are reported in Table 2.

## Methods

### Sample collection

Additional sequence data were generated for four ancient samples which were previously collected and described in the following original publications: ZVEJ25 and ZVEJ31 were published in Jones et al. (2017)^6^, KK1 in Jones et al. (2015)^7^ and NE5 in Gamba et al. (2014)^8^. Furthermore, 14 additional ancient samples and modern samples have been downloaded from the literature (see Table 1 and 2). The final dataset includes 34 samples consisting of 18 ancient and 16 modern samples.

**Table 1:** Metadata for ancient samples. Samples in bold have been resequenced in this study.

| Sample | Study | County | Site | Latitude | Longitude | Mean date BP | Date (2-sigma) | UDG-treated |
| --- | --- | --- | --- | --- | --- | --- | --- | --- |
| AHUR_2064 | Moreno-Mayar JV et al., 2018 | USA | Spirit Cave, Nevada | 37.41 | -122.08 | 10970 | 10770-11170 calBP | yes |
| Anzick-1 | Rasmussen M et al, 2014 | USA | Near Wilsall, Montana | 45.97 | -110.66 | 12632 | 12707–12556 calBP | no |
| Bichon | Jones et al. 2015 | Switzerland | Bichon | 47.1 | 6.87 | 13665 | 13560- 13770 cal BP | no |
| **KK1** | Jones et al. 2015 | Georgia | Kotias Klde | 42.25 | 43.27 | 9712 | 9529-9895 cal BP | yes |
| Kolyma1 | Sikora M et al, 2019 | Russia | Duvanni Yar | 68.6 | 159.1 | 9786 | 9668-9904 calBP | no |
| Loschbour | Lazaridis et al. 2014 | Luxembourg | Echternach | 49.81 | 6.4 | 8055 | 6220-5990 calBCE | yes |
| Mota | Gallego-Llorente M et al,2015 | Africa | Mota Cave, Gamo highlands of southwest Ethiopia | 6.80 | 38.17 | 4471 | 4524-4418 Cal BP | no |
| NE1 | Gamba et al. 2014 | Hungary | Polgar Ferenci hat | 47.88 | 21.19 | 7140 | 5310-5070 calBC | yes |
| **NE5** | Gamba et al. 2014 | Hungary | Kompolt-Kigyoser | 47.17 | 20.83 | 7050 | 5210-4990 calBC | yes |
| SF12 | Guenther et al. 2018 | Sweden | Stora Förvar, Sweden | 57.28 | 18 | 7700 | 7500-4000 cal BC | yes |
| Sumidouro5 | Sikora et al. 2017 | Brazil | Caverna do Sumidouro, Lagoa Santa, Brazil | -19.54 | -43.94 | 10391 | 10258-10524 (97.0%) calBP | no |
| sunghirIII | Moreno-Mayar JV et al., 2018 | Russia | Sunghir | 56.176 | 40.503 | 34093 | 35154-33031 calBP | yes |
| USR1 | Moreno-Mayar JV et al., 2018 | USA | Upward Sun River site (USR) | 64.98 | -150.54 | 11435 | 11600-11270 cal BP | yes |
| Ust_Ishim | Fu et al. 2014 | Russia | Ust'-Ishim, Omsk Oblast | 57.43 | 71.1 | 45000 | 45000 calBP (46880–43210 calBP at 95.4% probability) | yes |
| WC1 | Broushaki et al. 2016 | Iran | Wezmeh Cave | 34.05 | 46.59 | 9219 | 7455-7082 BCE | no |
| Yana1 | Sikora M et al, 2019 | Russia | Yana RHS | 70.43 | 135.25 | 31684 | 31321-32047 calBP | no |
| **ZVEJ25** | Jones et al., 2017 | Latvia | Zvejnieki | 57.78 | 25.24 | 7689 | 7791-7586 calBP | yes |
| **ZVEJ31** | Jones et al., 2017 | Latvia | Zvejnieki | 57.78 | 25.24 | 5965 | 6179-5750 calBP | yes |

**Table 2:** Metadata for modern samples. SGDP: Simon Genome Diversity Panel.

| Sample_ID | Sample_acronym | Population_ID | Country | Latitude | Longitude | Study |
| --- | --- | --- | --- | --- | --- | --- |
| SS6004477 | AUS | Australian | Australia | -13 | 143 | SGDP – Mallick et al., 2016 |
| LP6005443-DNA_B09 | DIN | Dinka | Sudan | 8.8 | 27.4 | SGDP – Mallick et al., 2016 |
| LP6005443-DNA_B03 | ESK | Eskimo_Sireniki | Russia | 64.4 | 173.9 | SGDP – Mallick et al., 2016 |
| LP6005519-DNA_D05 | IRU | Irula | India | 13.5 | 80 | SGDP – Mallick et al., 2016 |
| LP6005443-DNA_D04 | ITE | Itelman | Russia | 57 | 157 | SGDP – Mallick et al., 2016 |
| LP6005441-DNA_G06 | KAR | Karitiana | Brazil | -10 | -63 | SGDP – Mallick et al., 2016 |
| LP6005441-DNA_E07 | MND | Mandenka | Senegal | 12 | -12 | SGDP – Mallick et al., 2016 |
| LP6005443-DNA_G04 | MNS | Mansi | Russia | 63.65 | 62.1 | SGDP – Mallick et al., 2016 |
| LP6005441-DNA_F09 | ORQ | Oroqen | China | 50.4 | 126.5 | SGDP – Mallick et al., 2016 |
| LP6005443-DNA_D08 | PAP | Papuan | PapuaNewGuinea | -4 | 143 | SGDP – Mallick et al., 2016 |
| LP6005441-DNA_F10 | PIM | Pima | Mexico | 29 | -108 | SGDP – Mallick et al., 2016 |
| LP6005442-DNA_H12 | ULC | Ulchi | Russia | 52.43 | 140.42 | SGDP – Mallick et al., 2016 |
| LP6005442-DNA_D01 | XIB | Xibo | China | 43.5 | 81.5 | SGDP – Mallick et al., 2016 |
| LP6005442-DNA_F01 | YKT | Yakut | Russia | 63 | 129.5 | SGDP – Mallick et al., 2016 |
| LP6005442-DNA_B02 | YRI | Yoruba | Nigeria | 7.4 | 3.9 | SGDP – Mallick et al., 2016 |
| JHM06 | JHM | Jehai | Malaysia | 5.25 | 101.17 | McColl et al., 2018 |

### DNA extraction, Library preparation and next-generation sequencing

DNA was extracted and libraries were prepared for ZVEJ25, ZVEJ31, KK1 and NE5 (Table 3), following protocols described in the original publications, with the exception that DNA extracts were incubated with USER enzyme (5 μl enzyme: 16.50 μl of extract) for 3 hours at 37°C prior to library preparation in order to repair post-mortem molecular damage. The libraries were sequenced across 31 lanes of a HiSeq 2,500.

**Table 3:** Data statistics for newly sequenced samples. Average autosomal coverage was estimated on bam files after mapping quality filtering (mq20), duplicates removal, indel realignment and 2bp softclipping.

| Sample ID | Mass sampled (g) | Average autosomal coverage |
| --- | --- | --- |
| Kotias (KK1) | 0.101 | 12.03 |
| Latvia_HG2 (ZVEJ25) | 0.092 | 18.17 |
| NE5 (14.6) | 0.18 | 15.99 |
| ZVEJ31 | 0.102 | 9.97 |

### Bioinformatics analysis

#### Ancient samples

The following approach was used for both the newly sequenced ancient samples and the downloaded raw fastq files from previously published ancient samples.

Adapters were trimmed with Cutadapt v1.9.1^9^ and then raw reads were aligned to human reference sequence hg19/hs37d5 with bwa aln v0.7.12^10^ with seeding disabled (-l 1000), maximum edit distance set to -n 0.01 and maximum number of gap opens set to -o 2. Sai files were converted into sam files using bwa samse v0.7.12 and the read group line was also added. Bam files were generated using Samtools view v1.9^11^. Reads from multiple libraries belonging to the same sample were merged with the module MergeSamFiles within Picard v2.9.2^12^. Aligned reads were filtered for minimum mapping quality 20 with Samtools view v1.9. Indexing, sorting and duplicate removal (rmdup) were performed with Samtools v1.9. Indels were realigned using The Genome Analysis Toolkit v3.7^13^ (module RealignerTargetCreator and IndelRealigner) and 2bp were softclipped from the start and ends of reads using a custom python script. Final bam files were split by chromosome using Samtools view v1.9 and variant calling was performed with UnifiedGenotyper from The Genome Analysis Toolkit v3.7. All calls were filtered for minimum base quality 20 (-mbq 20) and reference-bias free priors were used (-inputPrior 0.0010 -inputPrior 0.4995). The same priors have been used for modern samples in the Simon Genome Diversity Panel^3^.

We focused on selecting a subset of the genome representing neutral genomic variation for demographic inferences^14, 15^. Therefore, specific filters were applied to discard: recombination hotspots (filter_hotspot1000g), poor mapping quality regions (filter_Map20), recent duplication (recent duplications, RepeatMasker score < 20), recent segmental duplication (filter_segDups), simple repeats (filter_simpleRepeat), gene exons together with 1000bp flanking and conserved elements together 100bp flanking (filter_selection_10000_100) and positions with systematic sequencing errors (filter_SysErrHCB and filter_SysErr.starch). All CpG sites were removed as well as C and G sites with an adjacent missing genotype. Genotypes were filtered by minimum coverage 8x and maximum coverage defined as twice the average coverage. Vcf files per chromosome belonging to the same sample were concatenated using vcf-concat from vcftools v0.1.15.^2 16^

#### Modern samples

Bam files were downloaded from the Simon Genome Diversity Panel^3^ and from McColl et al.^17^ (Table 2). Bam files were split by chromosome and variant calling, filtering for GC sites and coverage were performed as described above for the ancient samples with the same options and thresholds.

#### Final dataset

Per sample vcf files were compressed with bgzip and indexed with tabix from htslib v1.6^11^. The final dataset was assembled by merging filtered compressed vcf files for all modern and ancient samples with bcftools merge v1.6^11^. Only sites with called genotypes for all samples were kept using vcftools v0.1.15 (--max-missing 1). Triallelic sites were also discarded using bcftools view v1.6 (-m1 -M2). Final vcf statistics were generated with bcftools stats v1.6. Downstream analysis and plotting were performed in R v3.6.3^18^.

### Data Records

All newly generated sequencing raw reads have been deposited in the NCBI Sequence Read Archive XXX.

### Technical Validation

#### Summary of newly generated data

DNA was extracted for four previously published samples (ZVEJ25, ZVEJ31, KK1 and NE5) and sequence data were generated with an average coverage between 10x and 18x (Table 3). Endogenous DNA was estimated between 0.48 and 0.71 across all libraries (Table 4). Each library generated between 150 and 425 millions of reads corresponding to 15.2 and 42.9Gb respectively (Table 4).

**Table 4:** Raw data statistics for the newly sequenced libraries

| Sample | Total Bases | Read Count | GC (%) | Q20 (%) | Q30 (%) | Reads Aligned | Endogenous DNA |
| --- | --- | --- | --- | --- | --- | --- | --- |
| KK1_1 | 32,085,537,489 | 317,678,589 | 49.3 | 96.6 | 94.5 | 226,739,842 | 0.71 |
| KK1_2 | 31,821,488,543 | 315,064,243 | 49.7 | 96.9 | 94.8 | 221,241,435 | 0.70 |
| KK1_3 | 30,903,010,501 | 305,970,401 | 47.8 | 96.6 | 94.4 | 218,378,529 | 0.71 |
| KK1_4 | 28,374,056,452 | 280,931,252 | 48.5 | 96.6 | 94.5 | 200,616,589 | 0.71 |
| KK1_5 | 27,051,061,997 | 267,832,297 | 47.4 | 96.8 | 94.8 | 187,070,443 | 0.70 |
| KK1_6 | 26,428,490,321 | 261,668,221 | 49.7 | 96.7 | 94.5 | 182,602,757 | 0.70 |
| NE5_1 | 15,230,188,243 | 150,793,943 | 48.4 | 96.7 | 94.6 | 113,866,866 | 0.76 |
| NE5_2 | 22,443,822,868 | 222,216,068 | 47.8 | 96.7 | 94.6 | 167,444,317 | 0.75 |
| NE5_3 | 19,414,144,957 | 192,219,257 | 47.7 | 96.7 | 94.6 | 145,145,785 | 0.76 |
| NE5_4 | 35,602,627,361 | 352,501,261 | 48.9 | 96.8 | 94.7 | 257,297,424 | 0.73 |
| NE5_5 | 39,509,022,440 | 391,178,440 | 49.5 | 96.7 | 94.5 | 285,303,006 | 0.73 |
| NE5_6 | 38,119,633,918 | 377,422,118 | 47.7 | 96.8 | 94.7 | 275,284,926 | 0.73 |
| ZVEJ25_1 | 22,502,142,793 | 222,793,493 | 48.2 | 96.8 | 94.6 | 173,630,441 | 0.78 |
| ZVEJ25_2 | 26,264,479,451 | 260,044,351 | 47.5 | 96.8 | 94.6 | 202,756,810 | 0.78 |
| ZVEJ25_3 | 19,884,007,259 | 196,871,359 | 48.1 | 96.8 | 94.6 | 153,807,348 | 0.78 |
| ZVEJ25_4 | 30,314,118,184 | 300,139,784 | 47.0 | 96.9 | 94.8 | 234,102,091 | 0.78 |
| ZVEJ25_5 | 34,172,785,511 | 338,344,411 | 48.2 | 96.9 | 94.7 | 264,070,011 | 0.78 |
| ZVEJ25_6 | 32,515,172,804 | 321,932,404 | 48.2 | 96.9 | 94.7 | 251,187,453 | 0.78 |
| ZVEJ31_1 | 42,951,382,412 | 425,261,212 | 52.0 | 96.9 | 94.7 | 215,656,479 | 0.51 |
| ZVEJ31_2 | 41,717,115,447 | 413,040,747 | 50.7 | 96.9 | 94.8 | 209,910,986 | 0.51 |
| ZVEJ31_3 | 36,806,312,233 | 364,418,933 | 53.8 | 96.7 | 94.4 | 185,131,989 | 0.51 |
| ZVEJ31_4 | 34,986,764,509 | 346,403,609 | 51.3 | 96.9 | 94.6 | 166,115,737 | 0.48 |
| ZVEJ31_5 | 34,797,229,121 | 344,527,021 | 53.8 | 96.8 | 94.5 | 164,914,158 | 0.48 |
| ZVEJ31_6 | 39,275,860,102 | 388,869,902 | 52.0 | 96.8 | 94.6 | 185,999,314 | 0.48 |

#### Summary of the whole dataset including ancient and modern samples

The final dataset includes 34 samples with 509,348,047 sites in neutral regions before filtering (see Methods section for a detailed description of which regions were considered for variant calling). Sites not called across all samples (0% missing data allowed) were then discarded and 73,439,415 were retained. Multi-allelic sites (3811) were also removed bringing the final number of filtered sites to 73,435,604 (Table 5). Minimum and maximum coverage per sample within the final dataset is 11.3x and 55x respectively (within filtered intervals) with an average coverage across all samples of 30.1x (Table 5). We calculated the number of transitions (ts), transversions (tv) and the ts/tv ratio per sample (Table 5). As expected, all eight ancient samples that were not subjected to UDG-treatment showed a higher ts/tv ratio than their UDG-treated counterparts (see Figure 2), consistent with higher levels of DNA damage in these samples. The Brazialian sample Sumidouro 5 shows the highest excess of transition, possibly due to poor DNA preservation caused by environmental conditions. All other samples (both modern and UDG-treated ancient) showed similar ts/tv ratio with an average of 1.73, maximum and minimum of 1.76 and 1.63 respectively (see Table 5, Figure 2).

**Table 5:** variant calling summary per sample. DP: depth of coverage in filtered intervals for variant calling.

| Sample | Ref_Hom_sites | Alt_Hom_sites | Het_sites | Transitions (ts) | Transversions (tv) | Average_DP | ts/tv ratio |
| --- | --- | --- | --- | --- | --- | --- | --- |
| Xibo | 73380486 | 22850 | 32268 | 34876 | 20242 | 36.6 | 1.72 |
| Mansi | 73380645 | 21817 | 33142 | 34928 | 20031 | 45.6 | 1.74 |
| Oroqen | 73381419 | 23580 | 30605 | 34344 | 19841 | 39.0 | 1.73 |
| Ulchi | 73381180 | 23476 | 30948 | 34549 | 19875 | 42.0 | 1.74 |
| Yakut | 73380837 | 23102 | 31665 | 34610 | 20157 | 38.1 | 1.72 |
| Irula | 73379707 | 21860 | 34037 | 35402 | 20495 | 52.7 | 1.73 |
| Australian | 73380634 | 25423 | 29547 | 34826 | 20144 | 43.5 | 1.73 |
| Eskimo_Sireniki | 73382381 | 23785 | 29438 | 33827 | 19396 | 43.6 | 1.74 |
| Yoruba | 73366867 | 22452 | 46285 | 43520 | 25217 | 34.3 | 1.73 |
| Pima | 73383995 | 25261 | 26348 | 32647 | 18962 | 36.3 | 1.72 |
| Dinka | 73368528 | 22761 | 44315 | 42458 | 24618 | 36.0 | 1.72 |
| Karitiana | 73385473 | 25879 | 24252 | 31816 | 18315 | 44.2 | 1.74 |
| Mandenka | 73367366 | 22624 | 45614 | 43192 | 25046 | 33.2 | 1.72 |
| Papuan | 73381714 | 26484 | 27406 | 34211 | 19679 | 41.6 | 1.74 |
| Jehai | 73380775 | 23813 | 31016 | 34663 | 20166 | 36.0 | 1.72 |
| Itelman | 73382112 | 24509 | 28983 | 33903 | 19589 | 47.1 | 1.73 |
| SIII | 73380937 | 24070 | 30597 | 34878 | 19789 | 13.5 | 1.76 |
| kolyma1 | 73293180 | 24274 | 118150 | 120802 | 21622 | 16.3 | 5.59 |
| ahur_2064 | 73383950 | 24839 | 26815 | 32736 | 18918 | 20.0 | 1.73 |
| usr1 | 73382576 | 24728 | 28300 | 33352 | 19676 | 19.5 | 1.70 |
| yana1 | 73254076 | 23026 | 158502 | 161835 | 19693 | 28.8 | 8.22 |
| Bichon | 73244795 | 23656 | 167153 | 170509 | 20300 | 11.3 | 8.40 |
| WC1 | 73206319 | 21431 | 207854 | 209619 | 19666 | 11.9 | 10.66 |
| KK1 | 73381347 | 22877 | 31380 | 34269 | 19988 | 15.7 | 1.71 |
| Loschbour | 73383379 | 24998 | 27227 | 32383 | 19842 | 19.3 | 1.63 |
| ZVEJ25 | 73383085 | 23326 | 29193 | 33289 | 19230 | 23.2 | 1.73 |
| ZVEJ31 | 73381443 | 22542 | 31619 | 34352 | 19809 | 13.5 | 1.73 |
| mota | 73141456 | 23052 | 271096 | 269419 | 24729 | 13.6 | 10.89 |
| anzick-1 | 73014373 | 22982 | 398249 | 401458 | 19773 | 15.4 | 20.30 |
| NE5 | 73380544 | 21776 | 33284 | 34829 | 20231 | 20.8 | 1.72 |
| NE1 | 73193709 | 21302 | 220593 | 220990 | 20905 | 23.9 | 10.57 |
| Ust_Ishim | 73379574 | 21982 | 34048 | 35464 | 20566 | 35.2 | 1.72 |
| sf12 | 73383261 | 22971 | 29372 | 33185 | 19158 | 55.0 | 1.73 |
| sumidouro5 | 72439087 | 21290 | 975227 | 978128 | 18389 | 16.2 | 53.19 |

**Figure 2:**
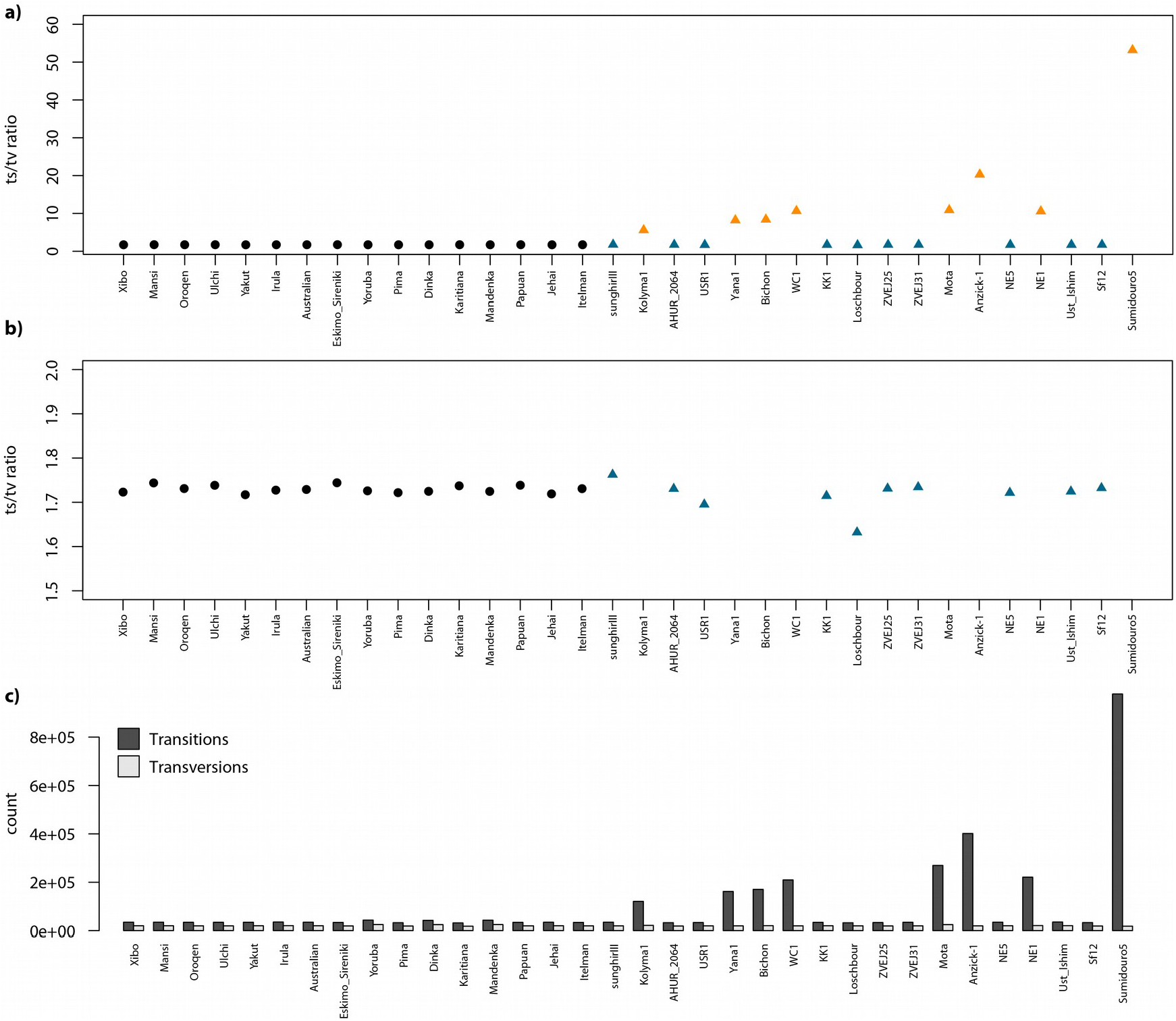
a) Transitions/Transversions ratio (ts/tv) per sample. Ancient and modern samples are represented by triangles and circles respectively. UDG and non-UDG treated samples are in blue and orange respectively. b) same as in a) but with a different y axis to focus on the ts/tv ratio among modern and UDG-treated ancient samples. c) Number of transitions (ts) and transversions (tv) per sample.

## Code Availability

The pipeline used to process the data with all scripts is available at XXX.

## Acknowledgements

PMD was supported by funding from the HERA Joint Research Programme “Uses of the Past” (CitiGen), the European Union’s Horizon 2020 research and innovation programme under Grant Agreement 649307. PMD and AM were supported by ERC Consolidator Grant 647797 ‘LocalAdaptation’. E.R.J. was supported by a Herchel Smith Research Fellowship. RP was supported by ERC starting grant ADNABIOARC (263441).

## Author contributions

AM designed the project. PMD, LC, EJ and AH performed the analyses. RP provided the samples. AM and PMD wrote the manuscript. All authors had input in the manuscript and approved the final version.

